# “Risk Factors for Visceral Leishmaniasis in Selected High Endemic Areas of Morang District, Nepal”: A case control study

**DOI:** 10.1101/530741

**Authors:** Punam Kumari Mandal, Rajendra Raj Wagle, Ajoy Kumar Thakur, Surendra Uranw

## Abstract

Visceral leishmaniasis is a major public health concern in Nepal. During the last few years, several KA outbreaks have been reported from Tarai region including Morang district. A case control study was conducted to assess the risk factors associated with VL in 5 endemic VDC of Morang district with 62 cases already treated from BPKIHS and Koshi zonal hospital and 248 controls selected randomly from the same village. Data collected using semi structured questionnaire from September to November 2013. This study revealed that people living in thatched house, sleeping in ground floor, ownership of animal, history of migration to India (Bihar and Jharkhand) and proximity to other KA cases within 50 m distance of household were strong risk factors for VL. Education remains protective (OR 0.39,95 % CI 0.19-0.79). The association with socioeconomic status showed clear dose – response effect. The odds for VL consistently decreased as the level of socioeconomic status increased (OR 4.26, 3.81). Strengthening surveillance system for early diagnosis and treatment, awareness programme and further extensive study is needed on risk factor, vector and control measures.

**Author Summary:** This study aims to explore the risk factors for visceral leishmaniasis. Based on findings there is a need to educate people in high-transmission areas how to realize, reduce or avoid environmental factors that favor the survival of the vector in the community. Similarly introduction of more exact surveillance tools in order to improve morbidity and mortality surveillance by health sector. People living in mud thatched houses need to be aware about cracks and crevices in the mud walls, their function as breeding places and how they can be controlled, for example by plastering with lime and mud․. However, a primary condition is that people need to understand the purpose of all these efforts in order to be motivated to put them into practice.

## Introduction

Kala-azar is known as a disease of the poorest of the poor because it predominantly affects the poor segment of the rural population with limited access to healthcare services [1]. In the endemic areas, children and young adults are its principal victims. Kala-azar is one of the major public health problem in Nepal with approximately eight million population are estimated at risk of disease i.e. 25% of the country population [4].

Kala-azar is first officially recorded in Nepal in 1980 from one district, Dhanusha [6]. So far, officially 12 districts in the central and eastern Terai (low lands) bordering to north Bihar state of India are endemic for KA. However, there is also increasing number of sporadic cases reported from other KA non-endemic districts including from the hilly regions [3]. During the last few years, several KA outbreaks have been reported from Terai region [6, 9]. Reporting of KA outbreak gradually has been increasing affecting many parts of the country including Morang district. Morang district is known to be Kala-azar endemic zone for long. There are a number of specific knowledge gaps that are particular interest to the elimination initiative in Nepal: Risk factor data are essential to design the appropriate public health response to an epidemic and purpose appropriate intervention to prevent future cases. Little is known about factors associated with Kala-azar in eastern Nepal and also have shown contradictory results related to the role of domestic animals as a risk factors in particular.

## Materials and Methods

The study design was case-control study. The objective was to explore the risk factors associated with Kala-azar at individual and households level in selected highly endemic areas of Morang districts, eastern Nepal.

### Study Population and Area

The study was conducted in Morang district. Morang district in the south close to the bordering district of Bihar state of India. There is open cross-border movement of people from Morang to Bihar state of India and vice versa due to sharing of similar culture and customs. Morang is administratively divided into 68 VDC and 9 VDC is the reportedly affected area of Kala-azar among which the study was conducted in 5 VDCs. These were Majhare, Bhathigach, Sisbanijahada, Katahari and Rangeli. These were the VDCs from where kala-Azar cases are reported relatively in higher number. The total no of cases from counted as 80 among these 62 cases were selected randomly.

### Ethics Statement

Before conducting the research approval was obtained from Institutional review board (IRB) and authorized letter was obtained from Department of Community Medicine and Public Health, Institute of medicine, Kathmandu for the purpose of data collection. Permission was taken from DPHO for medical records. The principle of human dignity and justice were maintained. An informed written consent was taken from each respondents, all adult patients and from a parent or guardian of participating minors before enrollment in the study. Patient’s medical records were reviewed retrospectively and all information retrieved from medical records was kept anonymized.

### Data Collection

Prior to preceding the data collection work, a sketch map of each sampled wards was prepared in consultation with the local key persons such as FCHVs, teachers, social workers etc. Then the list of households having kala-azar was prepared with the help of list available from DPHO Morang.

### Selection of Cases and Control

Kala-azar cases who were treated last 2 year to till date (January 2011 to till date) at Koshi zonal hospital and BPKIHS Dharan from the affected area were selected as a case according to list available from Medical records at the District Public Health Office (DPHO). Controls were selected randomly from the updated 2012/13 voting register of Morang district at the ratio of 1:4. Control were the healthy individual selected from the same population who had no suffered from VL in the past and did not present with fever and/or splenomegaly on the day of survey. Control were frequency matched for ages to cases to allow for inclusion of children in the group of control in the same proportion as cases, randomly selected adult voter were replaced by child.

The sample size was calculated through Epi Info [7] with following values based on study [15]. Two sided Confidence level = 95%, Power (1-β) = 80%,Case control ratio = 1:4 Percentage of control exposed = 38,Odds ratio = 2.4372,Percent of cases with exposure = 59.9. The sample size was 310 i.e. Cases=62 and control= 48

The study was conducted between September and November 2013. The total duration of data collection was of 8 weeks. Data were collected by researcher herself. The filled questionnaires were thoroughly checked and edited before data entry. Firstly, medical records at the District Public Health Office (DPHO), Morang was explored to compile the list of all Kala-azar who were treated last 2 years to till date and cases who gave ‘Morang district’ as their residence at the time of admission than those person were traced at their homes. Cases were all the confirmed KA cases those were treated at Koshi Zonal Hospital, Biratnagar and BPKIHS Dharan hospital. Controls were selected randomly from the updated 2012/13 voting register of Morang district. The register was based on a census of all citizens carried out in 2012/2013 in preparation of the elections and included all recent settlers at that time. To allow for inclusion of children in the group of controls in the same proportion as cases researcher replaced the randomly selected adult voter by a child. In such case instead of interviewing the adult voter, we enrolled child. If there were no children in the household sampled, a child was selected from the house of the nearest neighbor. A random sampling technique was used to enroll the controls from the village population using updated voting register of the affected area.

Data regarding household and risk factor were collected using pre-tested semi-structured questionnaire by principal investigator after obtaining the informed consent. Cases were asked to report their status at the time of their illness, for controls the status at the time of interview was recorded. In case of children, parents or guardians were interviewed.

Data was entered in Epi data software version 3.1 by investigator and analyzed SPSS Version 16. Descriptive analysis e.g. rate, confidence interval, medians with inter quartile ranges as required were calculated. The probability of the difference between cases and controls occurring by chance was tested by means of chi-square test. Risk factors were estimated by calculating the odds ratio (OR) as an approximation of the relative risk with 95% confidence interval (CIs). Observed associations were assessed through multivariate logistic regression. All variables with a P-value ≤ 0.10 in bivariate analysis were included in the multivariate logistic regression model after testing collinearity (tolerance >0.1 or VIF <10). Variables for the final model were selected using the backward elimination strategy. The probability of removal was set at *P* = 0.05.

Socio-economic status was assessed for the household, based validated asset index of NDHS. The asset index was converted into assets scores, using principle component analysis. Based on the asset scores, households were divided into five socioeconomic layers.

## Results

This study showed that majority of cases (29.0%) were from 13-25 years age group, males are more than females and indigenous caste (74.2%) were affected more than other caste. The majority of respondents were Hindu in comparison to others which comprises of 79% in cases and 81.5% in control where as other religion (Christian, Budhist, kirat) comprises 21% in cases and 18.5%. Majority of respondents were indigenous caste(Santhal+Mushar) by ethnic group being 74.2% cases and 68.5% controls followed by others which include Brahmin, chetri, baishya, janajati, dalitetc contributes 25.8 % in cases and 30.3% in controls. Similarly 31.1% cases and 60.5 % controls were literate, 35.5 % cases and 16.1% controls were from very low socioeconomic status.

Regarding the education the odds of contracting kala-azar in literate were 62% less than illiterate (OR 0.38 CI 0.21 - 0.68). Similarly in daily wage earner the risk of kala-azar is 2.15 times higher than others(student, preschool, housewife, private employee) (95% CI1.11 – 4.16). This study showed that respondents who were poor were more likely to develop VL than other. With thatched house, for those living in thatched house were 4.72 times higher risk of contracting the disease than brick house respectively (95% CI 2.29-9.71). The risk of having kala-azar was 2.42 times higher among those respondents whose house was nearby water collection (95% CI 1.36-4.29). Majority of cases 72.6% sleeps on the ground in comparison with controls 38.3%. The respondents who were sleeping on the ground were 4.26 times higher risk of getting the disease than the respondents who sleeps on bed (95% CI 2.31-7.88). Similarly, respondents who gave history of migration to India were 3.35 times higher risk of getting the disease than others (95% CI 1.84-6.10). The odds of having VL is 2.23 times higher in respondents who had relatives in Kala-azar endemic areas in India (95% CI 1.25-3.96). Whereas Ownership of bed net and alcohol use were not found significantly associated with occurrence of VL. This study revealed that people living in thatched house, sleeping in ground floor, ownership of animal, history of migration to India (Bihar and Jharkhand) and proximity to other KA cases within 50 m distance of household were strong risk factors for VL. Education remains protective (OR0.39,95 % CI0.19-0.79). The odds of getting VL among the respondents with proximity to VL cases within 50m distance was 2.63 times (95% CI 1.25-5.53).

The association with socioeconomic status remained significant and there was clear dose – response effect. The odds for VL consistently decreased as the level of socioeconomic status increased (OR4.26, 3.81).

## Discussion

This study shows the age group 13-25 years have higher number which comprises of 29% in both cases and control while in other studies conducted in national level and regional level shows incidence is very high among age group 15-40 years i.e. 60% [26]. It indicates that this age group has highest incidence of VL in comparison to other age group.

In this study the rate of infection was higher in indigenous caste than other caste i.e. 74.2% which suggest that indigenous caste have high chance of contracting the disease than others. Indigenous caste in this study comprises Santhal and Mushar. A study conducted on India suggest that VL typically clusters in marginalized communities of the villages at hamlet level [31], such as the mushar community in India [32]. Mushars (the lowest caste in Bihar) had twice the odds to be ‘late presenters’ compared to rest of castes (OR 2.05, 95% CI 1.24-2.38) [23]. In context of Nepal, more indigenous people (Santhal+ Mushar) resides in eastern region[36], this could be one of the reason that kala-azar is more endemic in eastern region than western region. Further study is recommended to verify this in future.

The higher rate of infection rate was found in men than women and similar ratio has been observed in relation to VL cases [10]. Although this differences is generally attributed to a more frequent exposure of males than females to sand flies,e.g as men often spend days away from home for seasonal work in farms, there could also be an under detection of disease in women in traditionally male dominated societies. Another behavioral-related factor is that women in those communities wear long dresses which could prove to be protective to some degree from sand fly biting.

Regarding the religion, Hindu were predominantly more than others. Literacy is a protective factor (OR 0.39, 95% CI 0.18-0.78). As expected VL cases comes from household with lower socioeconomic status than controls, compared with controls VL cases had thatched house, owned less land and more likely to be daily laborer. Kala-azar is related to poverty, affecting ‘the poorest of the poor’. In poor states such as Bihar in India and Nepal, VL affects families in the lowest income groups, who already live on less than US $ 1 per day. The relationship between leishmaniasis and poverty is complex: while poverty increases the risk for VL and aggravates disease progression, VL itself leads to further impoverishment of the family due to catastrophic health expenditure, income loss and death of wage earners [30].

The final model showed a very strong association between VL and certain housing factors, those living in a thatched house having 4.57 times higher odds of kala-azar. Sleeping on ground floor is a strong risk factor for kala-azar (OR 3.90, 95% CI 1.83-8.31). A number of studies in the Indian subcontinent investigated risk factors for VL, and most of them have been recently reviewed in detail by Bern et al. (2010) generally risk factors for VL were mainly linked to precarious housing conditions.VL is a rural disease and associated with precarious housing conditions (mud plastered house) and environment (humid soil and organic debris) of the poor communities. The proliferation of vector is enhanced by poor housing conditions which provide excellent breeding sites for sandflies and increased the risk of infection through the bite of vector or increased human-vector contact [7]. In this study concerning the association of living in thatched house with VL are supported by other studies in India that cracked house walls were associated with an increased risk of VL infection. A recent study fromIndia has shown evidence for mud plastered house as a risk factor for KA [14].this may happen in the mud walls of the human habitats or animal shelters. As high soil humidity favors an ideal breeding habitat for sand flies, they may be more attracted to house near water collection [13]. In this study presence of water collection nearby house was significant in bivariate analysis with odds of 2.42 (95% CI1.36-4.29). In rural areas, houses are usually surrounded by moderate-to-high density vegetation such as seasonal crops, bananas, bamboo trees, climbers and herbs. However, the presence of vegetation was not significantly associated in bivariate analysis.

In this study Proximity to previous kala-azar cases is also a risk factors for VL (OR 2.63, 95% CI 1.25-5.53) as 72.6% of cases wereliving in proximity of another kala-azar cases at 50m distance compared to 43.2% control. This clustering pattern suggests that active VL cases are the main source of transmission and is an important risk factor for l. donovani infection. The recent initiatives on active case detection strategy have included in index case based approach. The index case based approach includes the search of new VL and PKDL cases among the household members through house to house visits around a house (radius of 50m or 100 households) of a recently diagnosed VL case. It is important to increase the awareness of communities about the disease and its control. Specific adopted messages of health education should be developed. Community participation is essential in order to maximize the effect of control strategies, including case detection and vector control.

Ownership of animal was strong risk factors as respondents who had animal in their house having 3.95 times higher risk of contracting the disease than others (95% CI 1.87-8.37). The subjects who kept animal inside their house having 2.36 times higher risk of getting the disease (95% CI 1.57-5.35) in bivariate analysis but it was not significant in multivariate model. The result is same for keeping animal in sleeping room which was significant in bivariate analysis where as in multivariate model no association was observed. The role of domestic animals as a risk factor for VL is still controversial. The presence of cattle is associated with increased risk in some studies and decreased risk in others, reflecting the complexity of the effect of bovines on sand fly abundance, aggregation, feeding behavior and leishmanial infection rates [15]. In contrast to Latin America and Europe, where the host reservoir of VL is the domestic dog, humans are assumed to be the only reservoir on the Indian Subcontinent. Yet, domestic animals can play a role in the transmission of VL on the Indian subcontinent because of their association with the sandfly vector. Animals may either attract sandflies, thereby increasing vector density and transmission to humans; or they may serve as an alternative blood meal source, thereby decreasing transmission [11].

Similarly migration is a strong risk factor for VL those respondents who had history of migration to India (Bihar & Jharkhand) having 4.85 times higher odds of VL than others (95% CI 2.22-10.59). These findings reflect that these associations are all consistent with other studies as similar findings were found in a study conducted in Dharan [9].

### Bias and limitation

It was a retrospective study looking back at cases occurring over a period of 2 years for cases we asked for the conditions as they were at the time of their illness, for controls this was reflect the condition at present so there is likely to be recall bias. However, every possible effort was made during the interview to minimize the recall bias. Limitation of this study is the selection of controls, the controls were included for the study based on screening criteria not on the basis of laboratory investigation as serological test were not carried out to identify asymptomatic cases among the controls, which were selected primarily on the basis of the absence of clinical symptoms.

## Conclusion

Living in thatched house, ownership of animals, sleep on ground floor, history of migration to India (Bihar &Jharkhand), proximity to other KA cases within 50 m distance and poverty are the main risk factors associated with VL transmission in the VL endemic villages of Morang district. On the basis of the risk factor analysis, those who live in thatched house and sleeping on the ground appears to play an important role in the transmission of VL. Similarly, the ownership of animals also strongly associated with VL transmission.

There is a need to educate people in high-transmission areas of Morang district how to realize, reduce or avoid environmental factors that favor the survival of the vector in the community.

**Table 1.** Socio-demographic Characteristics of Cases and Controls (n=310)

| **Variables** | **Case(n=62)**<br>**(%)** | **Control**<br>**(n=248)**<br>**(%)** | **Total (n=310)**<br>**(%)** |
| --- | --- | --- | --- |
| Age (years) |  |  |  |
| 0-12 | 16(25.8) | 64(25.8) | 80(25.8) |
| 13-25 | 18(29.0) | 72(29.0) | 90(29.0) |
| 26-38 | 16(25.8) | 64(25.8) | 80(25.8) |
| 39+ | 12(19.4) | 48(19.4) | 60(19.4) |
| Median (inter quartile range) | 22.50(12-35) |  |  |
| Gender |  |  |  |
| Male | 35(56.5) | 106(42.7) | 141(45.5) |
| Female | 27(43.5) | 142(57.3) | 169(54.5) |
| Religion |  |  |  |
| Hindu | 49(79.0) | 202(81.5) | 251(81.0) |
| Others* | 13(21.0) | 46(18.5) | 59(19.0) |
| Ethnic group |  |  |  |
| Indigenousecaste** | 46(74.2) | 170(68.5) | 216(69.7) |
| Others | 16(25.8) | 78(31.5) | 94(30.3) |
| Education |  |  |  |
| Literate | 23(37.1) | 150(60.5) | 173(55.80) |
| Illiterate | 39(62.9) | 98(39.5) | 137(44.1) |
| Primary | 13(21.0) | 90(36.3) | 103(33.2) |
| Secondary | 8(12.9) | 55(22.2) | 63(20.3) |
| SLCand above | 2(3.2) | 5(2.0) | 7(2.3) |
| Occupation |  |  |  |
| Preschool | 6(9.7) | 21(8.5) | 27(8.7) |
| Student | 19(30.6) | 79(31.9) | 98(31.6) |
| Agriculture | 6(9.7) | 34(13.7) | 40(12.9) |
| Daily labour | 17(27.4) | 37(14.9) | 54(17.4) |
| Small scale business | 0(0.0) | 2(0.8) | 2(0.6) |
| Housewife | 13(21.0) | 73(23.4) | 86(27.7) |
| Private employee | 1(1.6) | 2(0.8) | 3(0.9) |

\*others include christian, buddhist and kirat; \*\* Indigenous caste include Santhal and Mushar

**Table 2.**
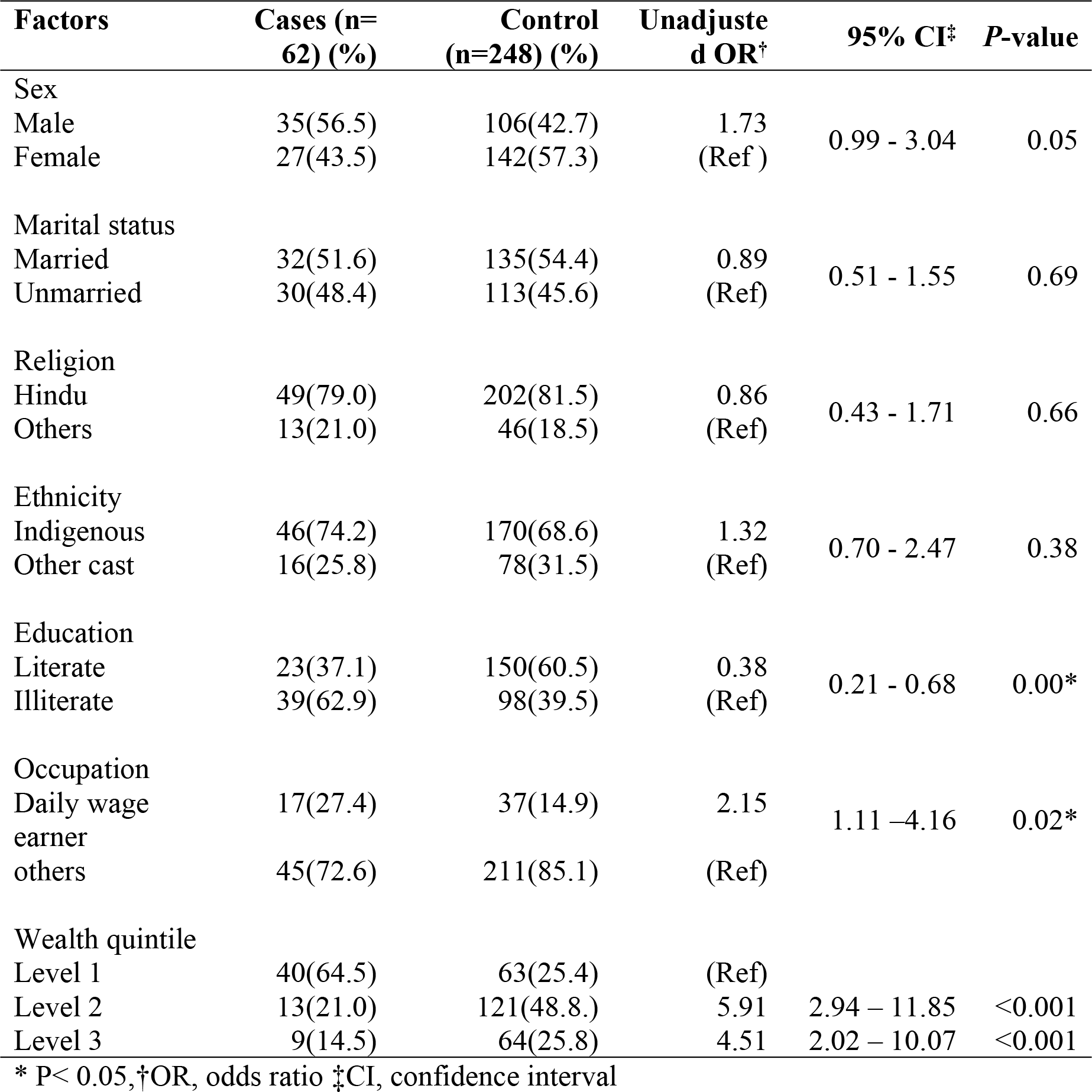
Socio-demographic and Economic Characteristics Associated with VL

**Table 3.**
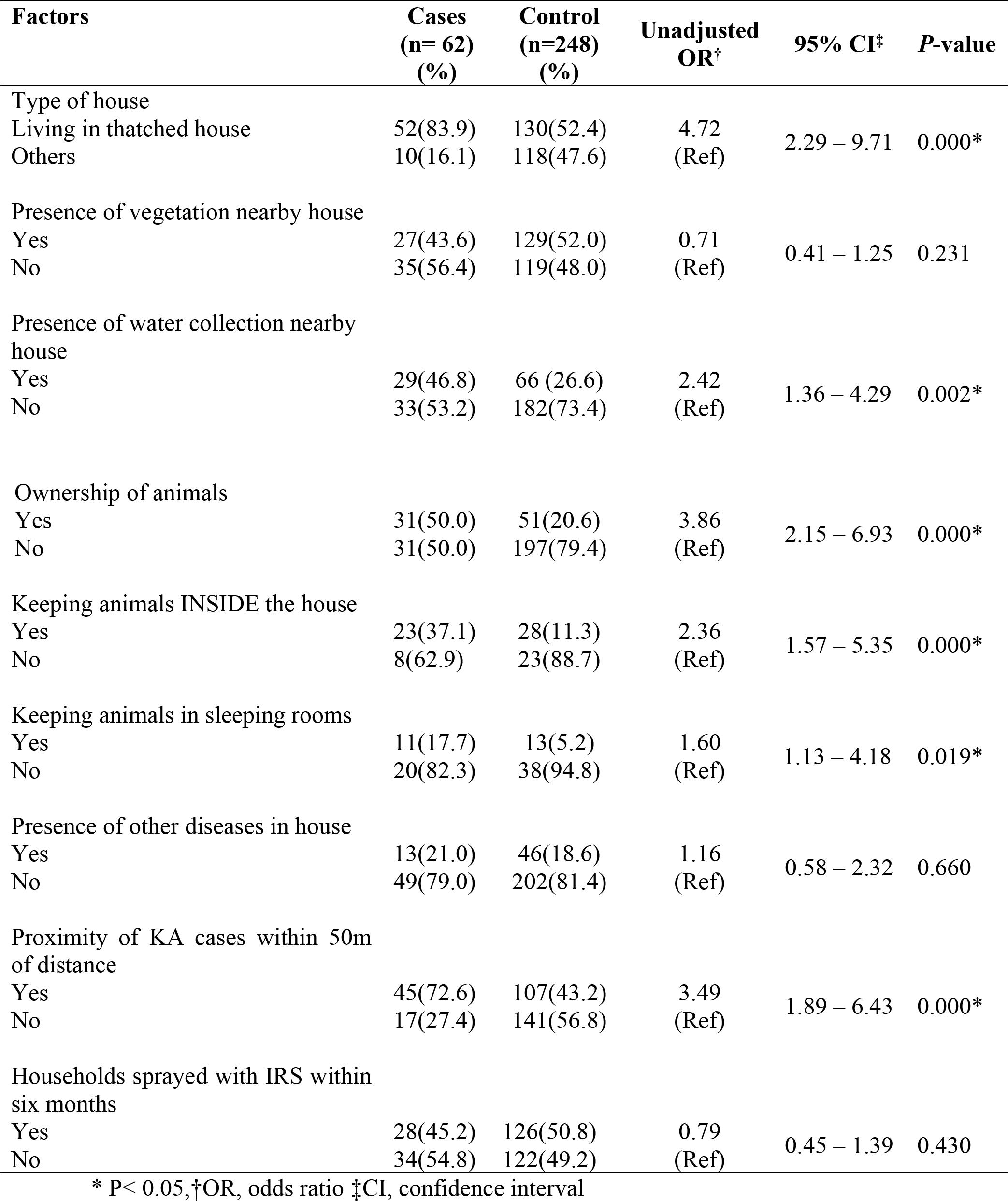
Household Factors Associated with VL

**Table 4.** Behavioral Factors Associated with VL

| **Factors** | **Cases**<br>**(n= 62)**<br>**(%)** | **Control**<br>**(n=248)**<br>**(%)** | **Unadjusted**<br>**OR<sup>†</sup>** | **95% CI<sup>‡</sup>** | **P-value** |
| --- | --- | --- | --- | --- | --- |
| Ownership of bed nets |  |  |  |  |  |
| Yes | 58(93.6) | 234(94.4) | 0.86 | 0.27 – 2.73 | 0.810 |
| No | 4(6.4) | 14(5.6) | (Ref) |  |  |
| Sleep under bed net last night |  |  |  |  |  |
| Yes | 52 (89.7) | 223(95.3) | 0.43 | 0.15 – 1.21 | 0.100 |
| No | 10(10.3) | 25(4.7) | (Ref) |  |  |
| Use of medicated bed nets |  |  |  |  |  |
| Yes | 21(36.8) | 23(10.0) | 5.22 | 2.62 - 10.41 | 0.000* |
| No | 41(63.2) | 225(90) | (Ref) |  |  |
| Sleeping habit |  |  |  |  |  |
| Sleep on the ground<br>on bed | 45(72.6)<br>17(27.4) | 95(38.3)<br>153(61.7) | 4.26<br>(Ref) | 2.31 – 7.88 | 0.000* |
| History of migration to India |  |  |  |  |  |
| Yes | 26(41.9) | 44(17.7) | 3.35 | 1.84 - 6.10 | 0.000* |
| No | 36(58.1) | 204(82.3) | (Ref) |  |  |
| Relatives in KA endemic areas<br>in INDIA |  |  |  |  |  |
| Yes | 39 (62.9) | 107 (43.2) | 2.23 | 1.25 – 3.96 | 0.005* |
| No | 23(37.1) | 141(56.8) | (Ref) |  |  |
| Alcohol use |  |  |  |  |  |
| Yes | 25(40.3) | 72(29.0) | 1.65 | 0.93 - 2.94 | 0.080 |
| No | 37(59.7) | 176(71.0) | (Ref) |  |  |

\* P< 0.05, <sup>†</sup>OR odds ratio, <sup>‡</sup>CI confidence interval

**Table 5.**
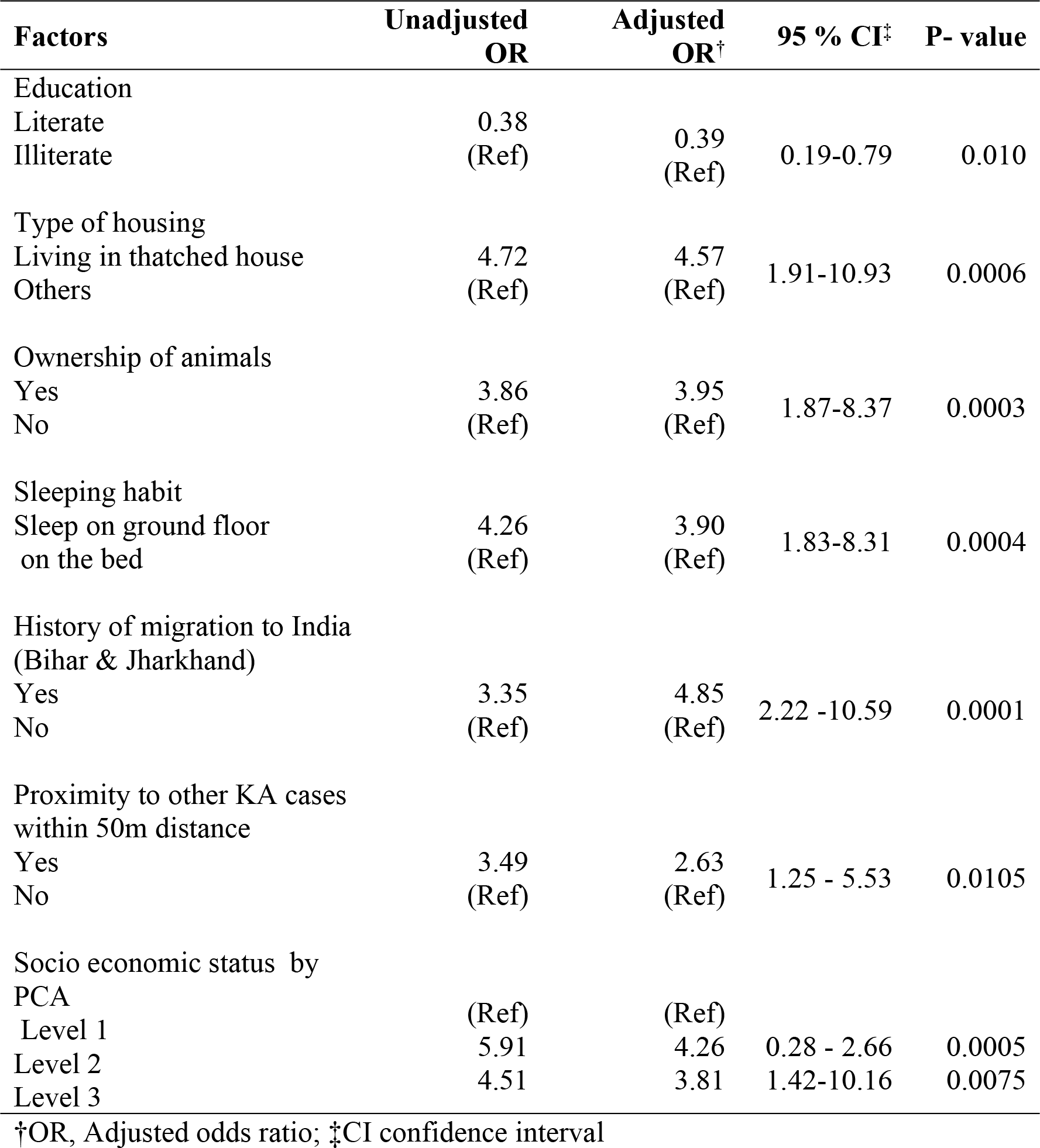
Factors Associated with VL in a ‘Multivariate’ Analysis

## Acknowledgement

I would like to express my sincere gratitude to Tribhuvan University,Institute of medicine for granting me opportunity to carry out this study. First of all I would like to express my sincere gratitude to Professor Dr.Rajendra Raj Wagle and Associate Professor Mr. Ajoy Kumar Thakur for continuous untiring guidance, valuable suggestion and inspiration. I am very much grateful to Dr. Surendra Uranw, Research coordinator BP Koirala institute of medical sciences for valuable guidance, supervision, suggestion and encouragement contributing in the completion of this study. I would like to extend my cardinal thanks to staffs of DPHO Morang. I am indebted to all the respondents for their kind co-operation and valuable time.

## Supporting information legends

**S1 Checklist: STROBE Checklist**

